# VaccineDesigner: A Web-based Tool for Streamlined Multi-epitope Vaccine Design

**DOI:** 10.1101/2024.03.20.585850

**Authors:** Dimitrios Trygoniaris, Anna Korda, Anastasia Paraskeva, Esmeralda Dushku, Georgios Tzimagiorgis, Minas Yiangou, Charalampos Kotzamanidis, Andigoni Malousi

## Abstract

**Background:** Multi-epitope vaccines have become the preferred strategy for protection against infectious diseases by integrating multiple MHC-restricted T-cell and B-cell epitopes that elicit both humoral and cellular immune responses against pathogens. Computational methods address various aspects independently, yet their orchestration is technically challenging as most bioinformatics tools are accessible through heterogeneous interfaces and lack interoperability features. The present work proposes a novel framework for rationalized multi-epitope vaccine design that streamlines end-to-end analyses through an integrated Web-based environment.

**Results:** VaccineDesigner is a comprehensive Web-based framework that streamlines the design of protective epitope-based vaccines by seamlessly integrating computational methods for B-cell, CTL, and HTL epitope prediction. VaccineDesigner incorporates single epitope prediction and evaluation as well as additional analyses, such as multi-epitope vaccine generation, estimation of population coverage, molecular mimicry, and proteasome cleavage. The functionalities are transparently integrated into a modular architecture, providing a single access point for rationalized, multi-epitope vaccine generation, time and cost-effectively.

**Conclusions:** VaccineDesigner is a Web-based interface for developing multi-epitope vaccines. Given a protein sequence, VaccineDesigner identifies and evaluates candidate B-cell, CTL, and HTL epitopes and constructs a library of multi-epitope vaccines that combine strong immunogenic responses, safety, and broad population coverage. The source code is available under the academic license at: https://github.com/BiolApps/VaccineDesigner. VaccineDesigner is freely accessible at: http://bioinformatics.med.auth.gr/VaccineDesigner.

## 1. Background

Developing effective and safe vaccines against pathogens is a leading research priority in immunology and public health. Vaccine development is typically based on live attenuated pathogens, inactivated agents, subunit formulations, or recombinant protein antigens. However, these methods raise several safety concerns, struggle with pathogen cultivation, and are time-consuming, therefore impractical particularly during pandemics^1^. During the last two decades, computational methods have become indispensable in trying to improve efficacy, safety, and population coverage, rationalizing vaccine design. Reverse Vaccinology (RV)^2^ is fundamentally based on computational methods to sift through the genetic information of a pathogen and identify efficient vaccine candidates that engage with the immune system. However, the identification of suitable proteins in pathogen proteomes is only the first step in computational vaccine design. Novel computational approaches have revolutionized our ability to predict and strategically harness epitopes^3^ i.e., tiny peptide fragments of antigens, capable of inducing protective immunity against pathogens. Since epitope-based approaches have been systematically used for some years now^4^ a reasonable advancement would be the identification of immunodominant epitopes in vaccine candidates and their combination in efficient multi-epitope peptide constructs.

Multi-epitope vaccines constitute a preferable strategy for several reasons, including the increased specificity over traditional methods, customizability, stability and convenient production^5^. In addition, multi-epitope vaccines can include different types of epitopes, epitopes of different antigens, epitopes of different pathogens for cross protection, or epitopes of different protein isoforms, for broader population coverage. On the antipode, multi-epitope vaccines are often inadequately immunogenic, and optimization is difficult as it is not feasible to exhaustively examine all possible peptide combinations^5^. Current methods on multi-epitope peptide construction are known from fusion protein technology^6^ and the strategic design of multi-epitope vaccines offers a promising approach for addressing diseases against a wide spectrum of pathogens, including parasites (visceral leishmaniasis^7^ and onchocerciasis^8^), viruses (COVID-19^9–10^ and Ebola^11^) and bacterial infections (Helicobacter pylori^12^ and Klebsiella pneumoniae^13^).

The components of a multi-epitope vaccine include B-cell epitopes for humoral responses (B-cell), cytotoxic T lymphocyte epitopes for cellular immunity (CTL), and helper T lymphocyte epitopes (HTL) for immune regulation^14^. B-cell epitopes consist of surface-accessible clusters of amino acids and activate immune responses through antibody production^15^. T-cell epitopes consist of small peptide fragments and activate immune responses by presentation to antigen-presenting cells (APCs) through MHC class I or class II molecules. Cytotoxic T-cells respond to MHC class I-restricted peptides, called CTL epitopes, whereas Helper T-cells target MHC class II-restricted peptides, referred to as HTL epitopes^16^. A multi-epitope vaccine has the unique ability to simultaneously activate both humoral and cellular immunity while avoiding the presence of elements that could provoke harmful reactions^17^. The final multi-epitope construct is important to include components with adjuvant capacity, to maximize immunogenicity as well as suitable linkers/spacers between epitopes, for optimal proteasome processing and minimization of junctional epitopes^18^.

Over the last decades, epitope-based computational approaches have revolutionized vaccine design. BepiPred 3.0^19^, BCEPRED^20^, and ABCpred^21^ rely on machine learning methods to predict linear B-cell epitopes, while CBTOPE^22^ integrates structural data to predict conformational epitopes. DiscoTope 3.0^23^ and ElliPro^24^ focus on the prediction of discontinuous B-cell epitopes based on the spatial arrangements of amino acids. To address T-cell epitope prediction, NetMHCpan-4.0^25^ and NetMHCIIpan-4.0^25^ quantify peptide-MHC binding affinities using a neural network architecture while PROPRED^26^ is primarily focused on identifying promiscuous MHC class-II binding regions. A wide range of tools have been developed to evaluate the properties of the predicted epitopes. ProtParam^27^, VaxiJen^28^, and Vaxign-ML^29^ assess the physicochemical characteristics, and the antigenic, protegenicity potential of the predicted epitopes, respectively. IFNepitope^30^ predicts IFN-γ inducing epitopes, while AlgPred 2.0^31^ and ToxinPred2^32^ assess the allergenic and toxic potential. Novel approaches attempt to combine outputs of multiple complementary tools in order to provide more effective and generalizable solutions. Vaxign2^33^, iVax^34^, and Vacceed^35^ are examples of workflows facilitating the discovery of vaccine targets in bacterial and eukaryotic pathogens. In support of these analyses The Immune Epitope Database and Analysis Resource (IEDB-AR)^36^ provides a robust and particularly valuable repository and analysis tool for experimentally validated immune epitopes and predictions.

Despite the advances in building integrated software tools^37^, the orchestration of RV resources remains technically challenging^38^ due to their structural heterogeneities and lack of interoperability^39^. Furthermore, existing frameworks have functional and technical limitations in terms of their ability to address epitope prediction for all cell types, ability to construct multi-epitope vaccines, and ease of use. To cope with these issues, we introduce a novel Web-based framework for rationalized multi-epitope vaccine design called VaccineDesigner. Rather than combining the outputs of individual tools, VaccineDesigner integrates and seamlessly execute cascading tasks for B-cell, CTL, and HTL epitope prediction through a user-friendly and fully customizable graphical interface. VaccineDesigner incorporates important functionalities such as multi-epitope vaccine generation, prioritization, population coverage estimation, molecular mimicry, and proteasome cleavage. The overall goal is to provide a comprehensive tool that supports end-to-end analyses, facilitating the prioritization of immunogenic epitopes into multi-epitope constructs likely to elicit protective immune responses.

## 2. Implementation

VaccineDesigner supports epitope prediction for B-cell and T-cell lymphocytes, as crucial indicators of protein antigenicity^40^. The architectural components are split into two discrete modules (Fig. 1). Module A *(Epitope prediction)* includes a set of tools for B-cell, CTL, and HTL epitope prediction, coupled with antigenicity, toxicity and allergenicity assessment for the candidate epitopes of all cell types. Module B *(Multi-epitope vaccine construction)* gets the individual tool predictions of Module A and builds a high-quality multi-epitope vaccine library that includes relevant vaccine components and quality filtering tools. VaccineDesigner applies the default parameter configuration of each tool in modules A and B, yet the parameters and thresholds of each tool are configurable according to user preferences.

**Figure 1:**
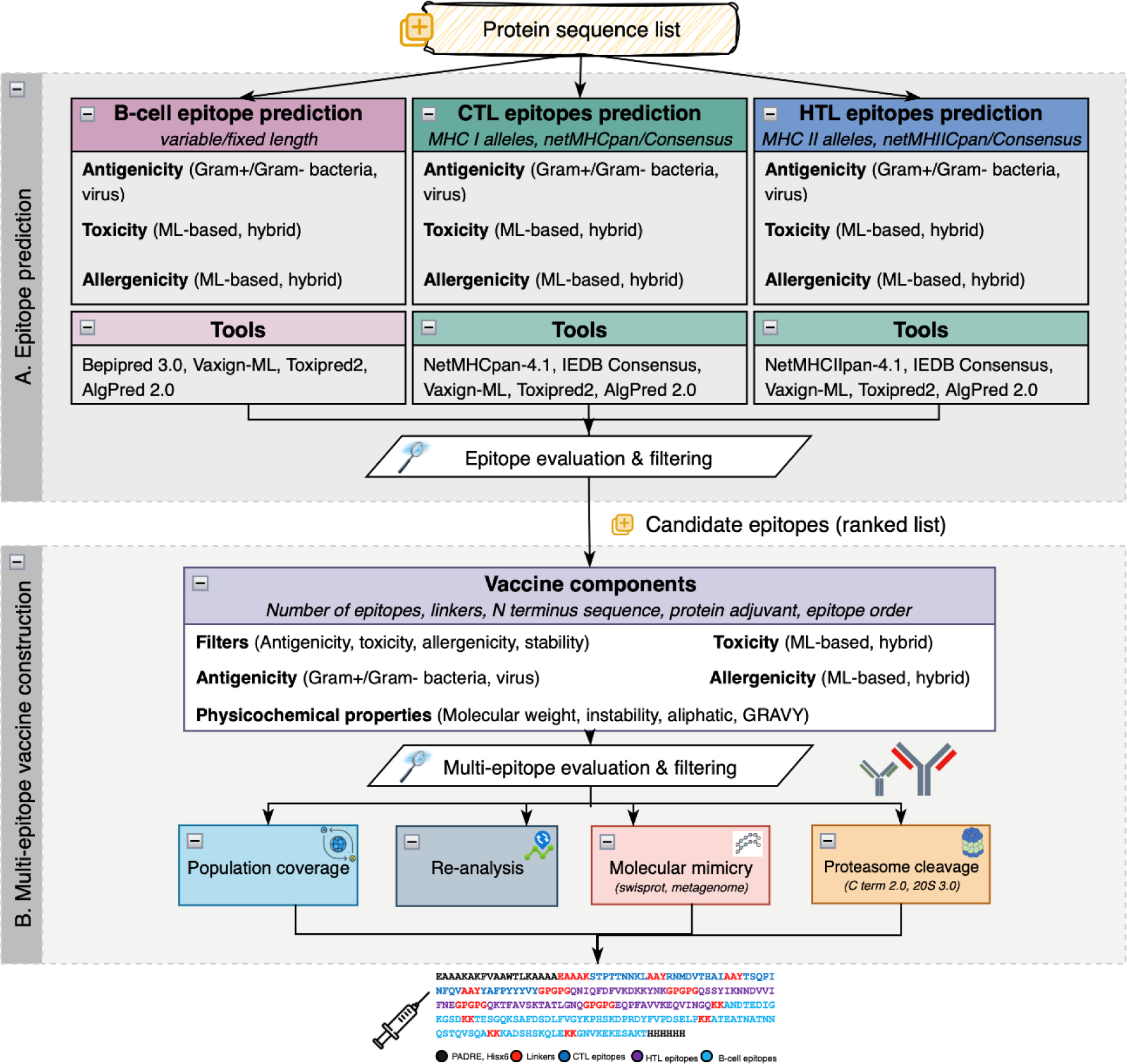
The main functional components of the streamlined process implemented by VaccineDesigner. **A.** The *Epitope prediction module* gets a fasta protein file, performs epitope prediction (B-cell, CTL, HTL), and ends up with the evaluation and filtering of epitope sequences, based on their antigenicity, toxicity and physicochemical properties. **B.** The *Multi-epitope sequence construction module* takes as input the tabular-formatted results containing the candidate epitopes and generates multi-epitope vaccine sequences based on user-defined parameters. To assess the quality of the candidate vaccine sequences the users are able to investigate the population coverage, molecular mimicry, proteasome cleavages, and to re-analyse the construct using the supported single-epitope prediction modules.

### 2.1 Epitope Prediction and Evaluation

#### 2.1.1 B-cell epitope prediction

B-cell epitopes are predicted using BepiPred 3.0. BepiPred 3.0 systematically scans protein sequences and assesses key physicochemical properties and sequence patterns to identify regions that can induce robust antibody responses^19^. Based on the amino acid scores VaccineDesigner generates B-cell epitopes, adhering to user-defined epitope length parameters. Two alternative analyses can be applied. The first involves the formation of B-cell epitopes in high-scoring regions. The lower length limit and the maximum number of amino acids between the predicted epitopes can be modified, enabling the combination of individual epitopes into larger immunogenic fragments. The second option pertains to the B-cell epitope generation. Users can define a fixed-length scanning window to assess the scores of all amino acids. Eligible epitopes are set as continuous amino acid regions with a user-defined length, wherein each position must fulfill certain filtering criteria.

#### 2.1.2 CTL epitope prediction

VaccineDesigner performs CTL epitope prediction using NetMHCpan 4.1^25^ and IEDB Consensus Method^41^. NetMHCpan implements an algorithm that identifies high-affinity regions for MHC class I alleles. Both methods predict strong and weak binder counts and export various information for the associated MHC alleles. The IEDB Consensus method combines the results of multiple prediction methods into a single consensus outcome providing a more reliable assessment, as prediction algorithms may produce varying results due to differences in underlying models and assumptions. The final ranking of CTL epitopes is determined by the number of alleles interacting with each epitope.

#### 2.1.3 HTL epitope prediction

HTL epitope prediction relies on the NetMHCIIpan 4.0 framework. NetMHCIIpan 4.0 identifies regions characterized by strong binding affinity for MHC class II molecules^25^. The results, akin to CTL epitopes, include strong and weak binders counts, along with details of the MHC class II alleles. HTL epitope prediction can be performed by the IEDB Consensus method for class II binding prediction, employing the same framework as for the Class I module^41^. The ranking of the HTL epitopes is determined by the number of interacting alleles, similar to the approach used for CTL epitopes.

#### 2.1.4 Epitope evaluation

Given a set of candidate B-cell, CTL, and HTL epitopes, VaccineDesigner applies various quality metrics to filter out epitopes with suboptimal antigenicity and safety profiles, adhering to user-defined thresholds. Vaxign-ML^29^ is used to comparatively assess the level of antigenicity for each epitope, prioritizing those that are highly immunogenic. To cope with potential safety issues, ToxinPred2^32^ is integrated into Module A to identify toxins within epitope sequences, and AlgPred 2.0^31^ to minimize the risk of allergic reactions in vaccine recipients. The epitopes that meet high-quality criteria for antigenicity, toxicity, and allergenicity are considered more likely to improve the overall efficacy and safety of the candidate multi-epitope constructs.

### 2.2 Multi-epitope Vaccine Sequences

The synthesis of multi-epitope constructs is considered a favorable approach to mitigate junctional immunogenicity and efficient epitope separation and presentation to T-cell and B-cell receptors^14^. VaccineDesigner expands the analysis of candidate epitopes to craft multi-epitope vaccine sequences. Using the most prominent epitopes, VaccineDesigner fuses larger constructs by merging individual epitopes with appropriate linker sequences (default: GPGPG/HTL, AAY/CTL, KK/B-cell)^6,14,42–46^ in a two-step procedure as follows:

#### 2.2.1 Vaccine Sequences Construction

The multi-epitope vaccine construct is built by a variable, user-defined number of B-cell, CTL, and HTL epitopes, as well as linker sequences between the epitopes and the N-terminus sequence. A variety of protein adjuvants are available for the synthesis of the multi-epitope vaccine sequences^47–50^. In addition, the order of epitope combination can be defined in arrangements of B-cell, CTL, and HTL epitopes, exemplified as B-C-H (B-cell - CTL - HTL). VaccineDesigner generates candidate vaccine sequences, that form a versatile library of epitope sequence combinations. Researchers can then explore diverse epitope combinations containing up to five epitopes from each category (B-cell, CTL, and HTL).

#### 2.2.2 Vaccine Candidate Sequences Evaluation and Selection

In the final step, the constructed candidate vaccine sequence library undergoes rigorous evaluation. The algorithm assesses each sequence against predefined filters, including Vaxign-ML for antigenicity, ToxinPred2 for toxin prediction, Algpred 2.0 for allergenicity, and ProtParam for physicochemical sequence analysis and stability^27^. User-defined parameters, such as the desired number of vaccine sequences are used to keep track of the number of candidates that meet all quality criteria. The algorithm exports the final multi-epitope sequence constructs when the preferred number of candidates is reached. This dynamic approach ensures the selection of vaccine sequences that meet strict measures for immunogenicity, safety, and biochemical attributes, offering the most promising candidates for further development and validation.

The candidate multi-epitope library is short-listed based on the ranking of the antigenicity, allergenicity, stability, and physicochemical properties of each construct. In addition, each candidate multi-epitope vaccine is scored by the deep neural network algorithm employed by DeepVacPred^51^. The analysis exports the list of candidate multi-epitope sequences once the user-defined maximum number of candidates is reached. The final candidate sequences are ranked based on the weighted sum of individual antigenicity, toxicity, allergenicity, stability, and the DeepVacPred rankings as follows: Let *S* be the vector of scores for each parameter, with *S*_*i*_ ∈ ℝ and *R*(*S*) the corresponding rankings with *R*(*S*_*i*_) ∈ ℤ and *R*(*S*) being a permutation of rankings, such that higher scores receive lower positions. In addition, let *W* be a vector of user-defined weights with *W*_*i*_ ∈ ℝ. The best-scoring multi-epitope construct is the one that yields the smallest possible sum of weighted rankings, across the *n* ranking criteria i.e.,

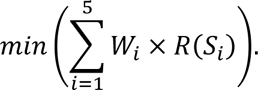

The newly generated sequences can be re-analyzed to verify the favorable properties of individual epitopes, in the whole construct, and can be further validated for population coverage, molecular mimicry, and the presence of putative proteasome cleavages. Population coverage is a highly recommended analysis for human vaccines and is implemented by the IEDB Population Coverage^36^ algorithm. To address molecular mimicry VaccineDesigner conducts protein similarity searches against SWISS-PROT^52^ or metagenomic proteins (env_nr) using the BLASTp^53^ algorithm. The analysis estimates whether a vaccine sequence shares similar domains with a known host protein, thereby ensuring its antigenicity and minimizing the risk of autoimmune reactions. Furthermore, proteasome cleavages can be identified on the protein sequences using NetChop 3.1^54^. The results are provided in both graphical and tabular formats as shown in Figure 2A.

**Figure 2:**
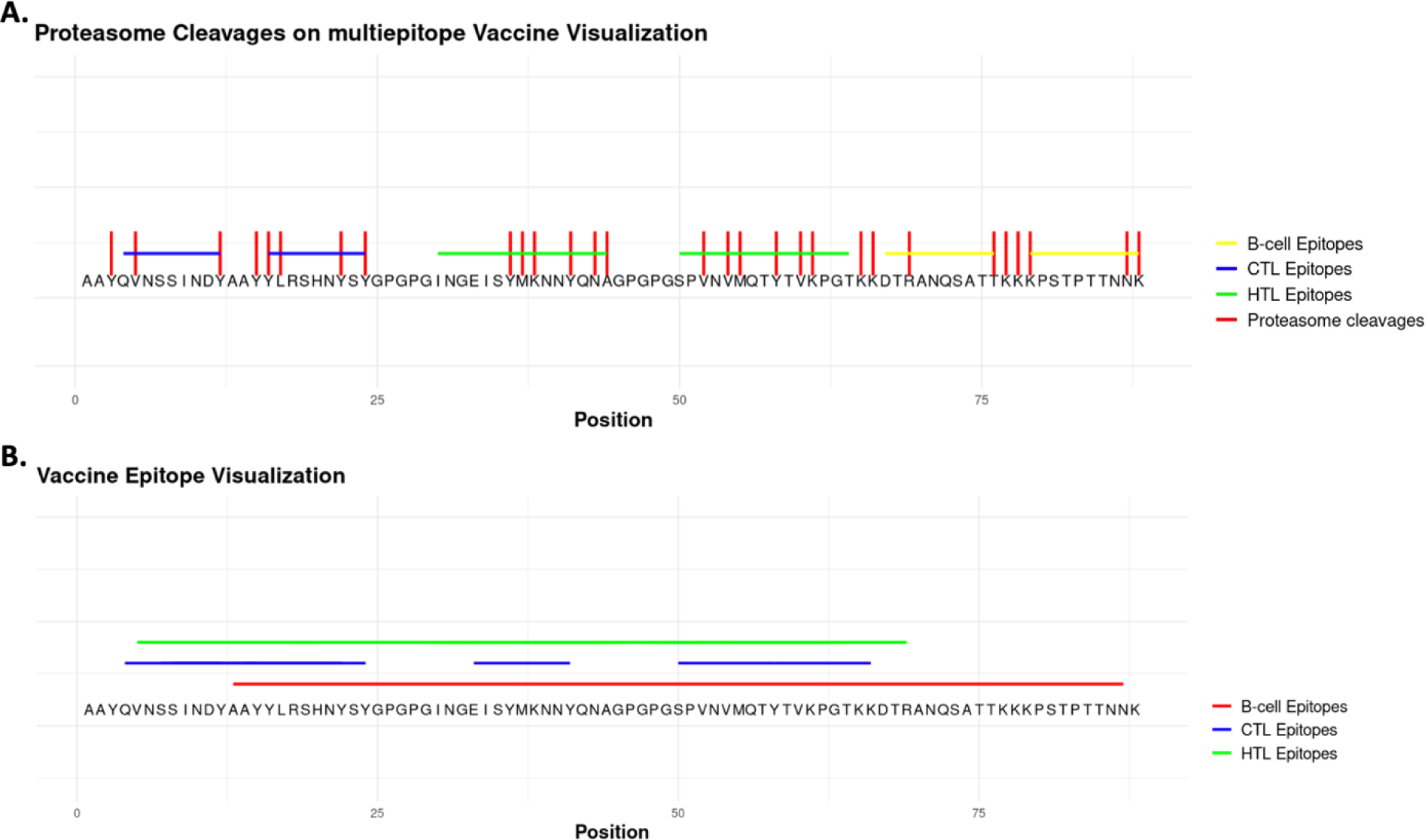
Visualization of multi-epitope vaccine based on the proteasome cleavage functionality (A), and multi- epitope sequence re-evaluation with the epitope prediction module (B).

## 3. Results

### 3.1 Usage scenario

To demonstrate the functionalities of the overall workflow, we implemented an example usage scenario on the *Atl* (UniProt: P0C5Z8) and the *IsdA* (UniProt: Q7A152) proteins of *Staphylococcus aureus*. *Atl* is a bifunctional autolysin playing a crucial role in the physiological processes, virulence, and pathogenesis of *S. aureus*. Autolysins are promising vaccine targets as they combine favourable features such as surface exposure, immunogenicity, and increased conservation among different strains. Iron-regulated surface determinant protein A (*IsdA*) is a virulence factor that is involved in the acquisition of iron, bacterial growth, and adherence to host tissues. *IsdA* combines desired vaccine properties including surface exposure on the cell wall of *S. aureus* and induction of immune responses in infected individuals. *IsdA* together with other cell wall-anchored surface proteins has been reported to induce significant protective immunity in murine models^55^.

In the usage scenario, 123 B-cell epitopes were predicted on the *Atl* and 136 on the *IsdA* protein using a fixed sequence length of 10 amino acids. 16 of these epitopes were considered high quality based on their quantified toxicity, allergenicity, and antigenicity measures. 96 and 18 CTL epitopes were identified in *Atl* and *IsdA,* respectively. These were predicted as strong or weak binders to MHC class I alleles, including *HLA-A*01:01, HLA-A*01:02, HLA-A*01:03, HLA-A*01:04, HLA-A*01:06, HLA-A*01:07, HLA-A*01:08* and *HLA-A*01:09*. The predicted epitopes underwent toxicity, allergenicity, and antigenicity filtering, resulting in 13 candidates. 193 and 50 HTL epitopes were considered strong or weak binders to MHC class II alleles for the *Atl* and the *IsdA* protein respectively, including *HLA- DRB1*01:01, HLA-DBR1*01:02, HLA-DBR1*01:03, HLA-DBR1*01:04, HLA-DBR1*01:05, HLA-DBR1*01:06, HLA-DBR1*01:07* and *HLA-DBR1\**01:08. Allergenic and toxic epitopes accounted for a significant number of epitopes resulting in 23 high-quality HTL epitopes for both proteins.

The list of the top-scoring epitopes of each category (B-CTL-HTL) was further narrowed down to four to reduce the computational cost. The candidates combined in the C-H-B order (CTL, HTL, and B-cell). Two epitopes of each cell type were fused in the final constructs, resulting in 1,728 candidate vaccine sequences. *Table 1a* presents the alignments of the newly generated multi-epitope vaccine sequences. *Table 1b* contains a ranking list of the first five multi-epitope sequences that passed all thresholds based on the level of antigenicity, toxicity, and relative properties.

**Table 1:** (a) A conserved domain of 98 amino acids in the five high-scoring multi-epitope sequences (top) and the aligned variable region of the 46 amino acid residues. (b): Combined ranking of the five multi-epitope sequence candidates along with their predicted antigenicity, toxicity, allergenicity, and instability index.

| Combined ranking | Sequence identifier | Antigenicity<br>(thr=50) | Toxicity<br>(thr=0.6) | Allergenicity<br>(thr=0.3) | Instability index<br>(thr=30) |
| --- | --- | --- | --- | --- | --- |
| 1 | seq2 | 79.29 | 0.298 | 0.281 | 39.09 |
| 2 | seq3 | 80.33 | 0.304 | 0.291 | 35.83 |
| 3 | seq4 | 78.88 | 0.347 | 0.282 | 38.35 |
| 4 | seq5 | 80.01 | 0.345 | 0.283 | 37.91 |
| 5 | seq1 | 90.86 | 0.304 | 0.291 | 34.86 |

The final ranking of the high-quality, multi-epitope sequences (combined ranking) is defined by the collective sum of individual rankings. The best scoring sequence (seq2) including linker sequences is shown in Figure 2. Figure 2A shows the outcome of the proteasome cleavage analysis in which the predicted cleavages are plotted for all the detected epitopes. Figure 2B depicts the multi-epitope sequence and its newly predicted B-cell, CTL and HTL epitope regions after re-evaluating the synthesized construct in the multi-epitope prediction module.

### 3.2 Evaluation against experimentally validated epitopes

To assess the competence of VaccineDesigner we sought to compare the predictions of linear B-cell epitopes and, cytotoxic and helper T-cell epitopes of SARS-CoV-2 Spike glycoprotein (UniProt ID: P0DTC2) against experimentally validated sequences provided by the IEDB database.

*Testing procedure:* The configuration parameters of each tool were set using the integrated graphical environment of VaccineDesigner. To quantify the level of antigenicity, Vaxign-ML was configured for viruses^21^ and filtered based on the protegenicity of each epitope (>40%). Since viruses do not produce toxins and their pathogenicity arises from their infection and replication cycle^56^, ToxinPred2 can be omitted or applied with high thresholds for testing purposes. In this example, ToxinPred2 and AlgPred 2.0 thresholds were set to 0.8 and 0.45, respectively. BepiBred 3.0 was used to predict B-cell epitopes of variable length. The threshold was set to 0.1, lower than the default 0.1512 to identify a broader range of epitopes and subsequently narrow down the predictions either by applying more stringent thresholds or by ranking the peptides based on other properties. The parameter *Subthreshold Amino Acid Inclusion Count* was set to 1 to enable the synthesis of larger continuous epitope regions that are separated by a single low-score amino acid (*Secondary threshold*: 0.08). CTL and HTL epitopes were predicted on the most common HLA alleles with positive assays that were collected from the IEDB database (MHC I molecules: *HLA-A*02:01*, *HLA-A*02:03*, *HLA-A*02:02*, *HLA-A*68:02*, *H2-Kb*, *HLA-A*24:02*, *HLA- A*11:01*, *HLA-E*01:01*, and MHC II molecules: *HLA-DRA*01:01/DRB1*04:01*, *HLA-DRB1*15:01*, *HLA-DRB3*02:02*, *HLA-DQA1*01:02/DQB1*06:02*, *HLA-DRB1*07:01*, *HLA-DRB5*01:01*). VaccineDesigner was set to run netMHCpan-4.1 and netMHCIIpan-4.0 with peptide length of 9 and 15, respectively.

*Results:* VaccineDesigner exported seventeen variable-length B-cell epitopes of which seven were filtered out due to the unqualified toxicity and/or antigenicity levels (Suppl. file). All predictions overlap with on average 40 experimentally validated epitopes. The highest scoring B-cell epitope is predicted in a non-toxic and non-allergen 74 amino acids sequence that matches the region including the most positive assay counts. Figure 3A shows a significant overlap between the predicted B-cell epitopes with the experimentally validated ones. Similarly, of the 242 CTL and 346 HTL predicted epitopes we selected the top fifteen sequences with the highest number of strong and weak binders for comparison against validated IEDB epitopes (Suppl. file). Figures 3B and 3C show the residual overlaps between the predicted MHC-I and MHC-II epitopes and the validated 263 MHC-I and 1,783 MHC-II epitopes from the IEDB positive assays (IEDB v.3.10.0). Evidently, most CTL and HTL predictions are collocated with known epitopes. CTL predictions are on average more accurate compared to HTL which is possibly related to the significantly higher number of positive assays implying stronger experimental evidence (on average 11.4 CTL vs 1.6 positive assay counts). However, further comparative analyses are needed to assess how different parameter configurations can affect the overall performance which is not realistic to implement manually. By enabling transparent execution of the pipelined processes VaccineDesigner reduces the execution time which is particularly important when a candidate protein must be examined exhaustively under different configurations.

**Figure 3:**
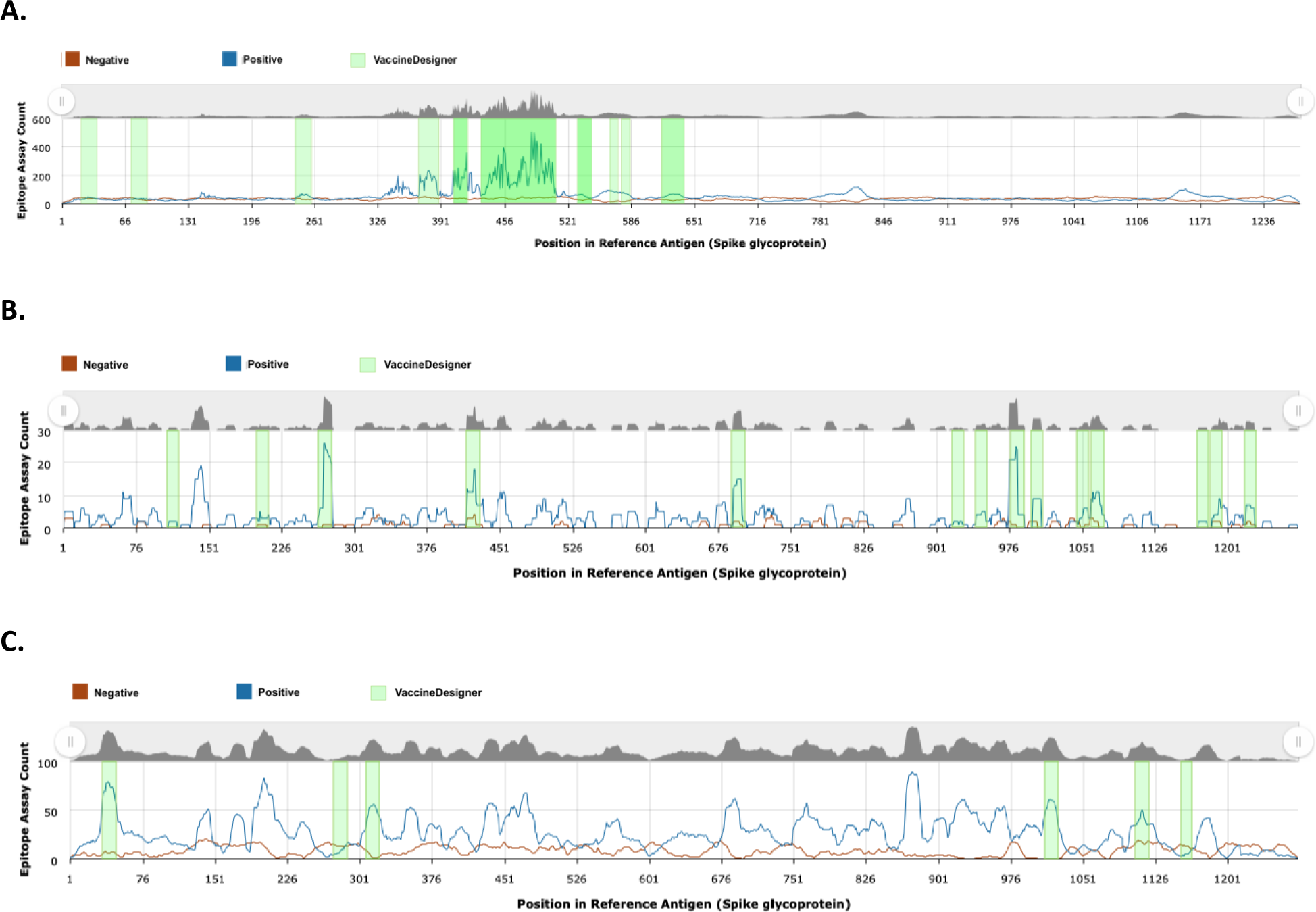
Overlapping residual loci between the top-scoring B-cell (A), CTL (B), and HTL (C) predictions and the experimentally validated positive and negative assays (IEDB v.3.10.0).

### 3.3 Comparison with other RV pipelines

During the last two decades, several RV pipelines have been developed. Table 2 lists the key attributes of the most popular RV implementations including the availability of graphical interfaces, streamlined processes, support for different epitope types, prediction capabilities for non-human species, multi-epitope sequence construction, pathogen protein selection, primary methods for vaccine candidate evaluation, and source code availability. Most of the RV tools predict epitopes following various approaches however, VaccineDesigner is the only tool that predicts and combines epitopes and then filters the multi-epitope constructs to select only the best epitope combinations. In addition, VaccineDesigner has the unique flexibility to enable modifications of the number of epitopes from each category and select epitopes from different proteins based on their quality and user preference.

**Table 2:** Main features of RV pipelines.

| Name/ Tool | Graphical User Interface (GUI) (Yes/No) | Streamlined process (Yes/No) | Prediction for different epitope types (BP/TP/BP+TP) | Epitope prediction for non-human species (Yes/No) | Multi-epitope sequence construction (Yes/No) | Pathogen protein selection (Yes/No) | Vaccine Candidate Evaluation (*) | Source code availability (Yes/No) |
| --- | --- | --- | --- | --- | --- | --- | --- | --- |
| NERVE<sup>57</sup> | No | Yes | No | No | No | Yes | SL, AP, HS | Yes |
| iVAX<sup>34</sup> | Yes | Yes | TP | Yes | Yes | Yes | I, C | No |
| OptiVax<sup>58</sup> | No | Yes | TP | Yes | Yes | Yes | I, PC, SP | Yes |
| ReVac<sup>59</sup> | No | Yes | BP + TP | Yes | No | Yes | A, C, SL, CR | Yes |
| IEDB-AR<sup>36</sup> | Yes | No | BP + TP | Yes | No | No | No | Yes |
| Vacceed<sup>35</sup> | No | Yes | TP | Yes | No | Yes | SL, SP, TH, T | Yes |
| VacSOL<sup>60</sup> | Yes | Yes | BP+TP | No | No | Yes | SL, V, TH, A, B+T | Yes |
| Vaxign2<sup>33</sup> | Yes | Yes | TP | No | No | Yes | SL, TH, AP, B&T | Yes |
| VaccineDesigner | No | Yes | BP+TP | Yes | Yes | No | B+T, A, CR, S, PC | Yes |

\***SL:** Subcellular Localization, **AP:** Adhesin Probability, **HS:** Host Similarity, **I:** Immunogenicity, **C:** Conservation, **PC:** Population Coverage, **A:** Antigenicity, **CR:** Cross Reactivity, **SP:** Signal Peptides, **TH:** Transmembrane Helices, **V:** Virulence, **BP:** B-cell prediction, **TP:** T-cell prediction, **BP+TP:** B-cell and T-cell prediction, **B+T:** Presence of B-cell and T-cell epitopes, **T:** Presence of T-cell epitopes, **S:** Stability

## 4. Discussion

VaccineDesigner was developed to build protective multi-epitope constructs using in silico B-cell, CTL, and HTL epitope prediction methods coupled with toxicity, allergenicity, and antigenicity assessment. Putative multi-epitope vaccines are further examined to estimate population coverage, molecular mimicry, and proteasome cleavage to maximize protective immune responses. Technically, VaccineDesigner aims to address the reported underutilization of automated workflows in reverse vaccinology by enabling pipelined analyses through a user-friendly graphical environment that includes preconfigured software tools and effortless I/O data flow management^61^.

VaccineDesigner extends RV frameworks by introducing several technical and functional advancements: 1) Implements a stepwise procedure that starts with a list of candidate antigenic proteins, and progressively leads to the most likely effective multi-epitope vaccines. 2) Leverages the collective strengths of each tool and combines their output in unified tabular and graphical reports. 3) Applies a weighted scoring scheme to prioritize multi-epitope constructs based on individual properties. 4) Ensures time-efficient and effortless application due to the seamless data flow between cascading tasks implemented by individual tools. 5) Is independent of local installations and operating system restrictions as it is available through a fully customizable Web-based interface.

Considering the architectural components, VaccineDesigner is built on a foundation of well-validated methodologies recognized and trusted within the scientific community. The selection of the tools was primarily based on benchmarking studies^29,61^, citation-based recognition by the scientific community, and also on the technical constraints that are related to the programmatic access and availability of the source code. A feasible enhancement enabled by the modular architecture of VaccineDesigner is to combine evidence from multiple tools performing the same analysis. However, the lack of interoperable tools puts additional technical challenges as it is hard to standardize inputs, parameters, and outputs and most importantly to provide sophisticated and more effective consensus algorithms. Considering the significant number of informative properties that are produced for each candidate protein, a reasonable enhancement would be to mine discriminative properties through machine learning models to predict peptides that induce protective immune responses. Still, the impact of these models is limited by the inaccuracies of the positive and negative training data and the lack of a definitive list of analyses that all RV pipelines should include^61^.

It should be noted that VaccineDesigner does not address the initial protein selection stage. Practically, the starting point of the pipeline is a set of proteins that have already been selected as potentially immunogenic. To incorporate this analysis several challenges must be addressed including the selection of proteins based οn the subcellular localization, the exclusion of homologs to avoid autoimmune reactions in the host, and the use of pangenomes to select core genome components that reduce the selective pressure^62^. Moving to more holistic approaches VaccineDesigner will pursue the integration of 3D modelling algorithms to predict the folded 3D structure of the multi-epitope vaccine constructs and to identify exposed peptides that are more likely to interact with immune cells. Alphafold 3 revolutionizes research in this field and is expected to accelerate research in precision vaccine discovery by predicting the structures of DNA, RNA, and molecules like ligands, which are essential to drug discovery^63^.

## 5. Conclusions

In the evolving landscape of infectious diseases, multi-epitope vaccine design has emerged as a pivotal strategy to identify antigenic determinants in pathogenic proteins, while also reducing the risk of adverse reactions and insufficient coverage against diverse pathogenic strains. VaccineDesigner is the only reverse vaccinology framework that combines an automated procedure for the prediction and assessment of protective multi-epitope constructs with a graphical and fully customizable Web-based environment. As our insights into the host-pathogen interactions advance, we anticipate that VaccineDesigner will provide a valuable tool for rationalized knowledge synthesis and will contribute to real-world applications as a generic framework for developing safer and more effective vaccines.

## Declarations

### Abbreviations

APCs: antigen-presenting cells
CTL: Cytotoxic T cells
HTL: Helper T cells
IEDB: Immune Epitope Database
IEDB-AR: Immune Epitope Database and Analysis Resource
MHC: Major Histocompatibility Complex
RV: Reverse Vaccinology

### Ethics approval and consent to participate

Not applicable

### Consent for publication

All authors gave their consent for publication

### Availability of data and materials

VaccineDesigner is an open-source tool freely available under the academic license at https://github.com/BiolApps/VaccineDesigner

Project name: VaccineDesigner

Project home page: http://bioinformatics.med.auth.gr/VaccineDesigner Operating system(s): Platform independent

Programming language: R, Python Other requirements: Web browser License: Academic Free License

Any restrictions to use by non-academics: license needed

### Authors’ contributions

A. T. contributed to: design of the work, creation of new software, preparation of the manuscript A.K. contributed to: design of the work, acquisition, analysis, preparation of the manuscript A.P. contributed to: acquisition, analysis, interpretation of data E.D. contributed to: acquisition, analysis, interpretation of data G.T. contributed to: interpretation of data, manuscript revision M.Y. contributed to: interpretation of data, manuscript revision C.K. contributed to: design of the work, acquisition, analysis, manuscript revision A.M. contributed to: conception, design of the work, creation of new software, preparation of the manuscript All authors have approved the submitted version and have agreed both to be personally accountable for the author’s own contributions and to ensure that questions related to the accuracy or integrity of any part of the work

## Acknowledgments

Not applicable

## Supplementary file

List of predicted B-cell, CTL, and HTL epitopes in Spike S glycoprotein (P0DTC2) of SARS-CoV-2 and the number of validated matching epitopes from IEDB database.

